# Evolution of increased larval competitive ability in *Drosophila melanogaster* without increased larval feeding rate

**DOI:** 10.1101/029249

**Authors:** Manaswini Sarangi, Archana Nagarajan, Snigdhadip Dey, Joy Bose, Amitabh Joshi

**Author notes:** equal contribution (Manaswini Sarangi, Archana Nagarajan). Email addresses.

## Abstract

Multiple experimental evolution studies on *D. melanogaster* in the 1980s and 1990s indicated that enhanced competitive ability evolved primarily through increased larval tolerance to nitrogenous wastes and increased larval feeding and foraging rate, at the cost of efficiency of food conversion to biomass, and this became the widely accepted view of how adaptation to larval crowding evolves in fruitflies. We recently showed that populations of *D. ananassae* and *D. n. nasuta* subjected to extreme larval crowding evolved greater competitive ability without evolving higher feeding rates, primarily through a combination of reduced larval duration, faster attainment of minimum critical size for pupation, greater efficiency of food conversion to biomass, increased pupation height and, perhaps, greater urea/ammonia tolerance. This was a very different suite of traits than that seen to evolve under similar selection in *D*. *melanogaster* and was closer to the expectations from the theory of *K*-selection. At that time, we suggested two possible reasons for the differences in the phenotypic correlates of greater competitive ability seen in the studies with *D. melanogaster* and the other two species. First, that *D. ananassae* and *D. n. nasuta* had a very different genetic architecture of traits affecting competitive ability compared to the long-term, laboratory populations of *D. melanogaster* used in the earlier studies, either because the populations of the former two species were relatively recently wild-caught, or by virtue of being different species. Second, that the different evolutionary trajectories in *D. ananassae* and *D. n. nasuta* versus *D. melanogaster* were a reflection of differences in the manner in which larval crowding was imposed in the two sets of selection experiments. The *D. melanogaster* studies used a higher absolute density of eggs per unit volume of food, and a substantially larger total volume of food, than the studies on *D. ananassae* and *D. n. nasuta*. Here, we show that long-term laboratory populations of *D. melanogaster*, descended from some of the populations used in the earlier studies, evolve essentially the same set of traits as the *D. ananassae* and *D. n. nasuta* crowding-adapted populations when subjected to a similar larval density at low absolute volumes of food. As in the case of *D. ananassae* and *D. n. nasuta*, and in stark contrast to earlier studies with *D. melanogaster*, these crowding-adapted populations of *D. melanogaster* did not evolve greater larval feeding rates as a correlate of increased competitive ability. The present results clearly suggest that the suite of phenotypes through which the evolution of greater competitive ability is achieved in fruitflies depends critically not just on larval density per unit volume of food, but also on the total amount of food available in the culture vials. We discuss these results in the context of an hypothesis about how larval density and the height of the food column in culture vials might interact to alter the fitness costs and benefits of increased larval feeding rates, thus resulting in different routes to the evolution of greater competitive ability, depending on the details of exactly how the larval crowding was implemented.

## Introduction

The theory of density-dependent natural selection is perhaps the most successful bridge between population genetics and population ecology, being premised on the notions that genotypic fitnesses are functions of population density, and that often no single genotype will be the most fit at both low and high densities (Mueller 2009). Following the early development of simple models of density-independent (*r*-selection) and density-dependent (*K*-selection) selection by MacArthur (1962) and MacArthur and Wilson (1967), a loose verbal theory attempting to explain life history variants via the *r*- and *K*-selection dichotomy took form (Pianka 1970). Subsequently, formal population genetic models of density dependent selection were developed (Gadgil and Bossert 1970; Roughgarden 1971; Clarke 1972; Asmussen 1983; Anderson and Arnold 1983), and these models, *inter alia*, highlighted the importance of *r*-*K* tradeoffs in mediating the evolutionary response to differences in population density. The first rigorous attempt to study *r*- and *K*-selection empirically was through laboratory selection on *Drosophila melanogaster* populations maintained either at low density by culling (*r*-populations), or at high density by allowing populations to reach their carrying capacity (*K*-populations). This work revealed a trade-off between *r* and *K* for the first time (Mueller and Ayala 1981): the *K*-populations had a higher per capita rate of population growth than the *r*-populations when assayed at high densities, but a lower per capita population growth rate than the *r*-populations when assayed at low densities. Further experiments showed the *K*-populations to have evolved a greater larval competitive ability (Mueller 1988), larval feeding rate (Joshi and Mueller 1988), foraging path length (Sokolowski et al. 1997), pupation height (Mueller and Sweet 1986, Joshi and Mueller 1993), but a reduced food to biomass conversion efficiency (Mueller 1990) such that larvae from the *K*-populations required more food to successfully complete development, as compared to the control *r*-populations. This finding, at odds with the classical expectation of the evolution of greater efficiency of food utilization under *K*-selection (MacArthur and Wilson 1967), along with the facts that the *K*-populations experienced higher densities than the *r*-populations both as larvae and adults, and that the *r*-and *K*-selection contrast was confounded with a discrete versus overlapping generation contrast, motivated a second selection study using *D. melanogaster* populations from a different geographic origin than the *r*- and *K*-populations, and selected for adaptation only to larval crowding (the CU populations, first described in Mueller et al. 1993).

Relative to the low larval density control populations (UU populations: Joshi and Mueller 1996), the CU populations evolved a set of traits very similar to that seen in the earlier study of the *K*-populations. The CU populations evolved greater pre-adult survivorship when assayed at high larval density (Mueller et al. 1993; Shiotsugu et al. 1997), largely through the evolution of increased larval feeding rate (Joshi and Mueller 1996), larval foraging path length (Sokolowski et al. 1997) and tolerance to nitrogenous wastes like urea (Shiotsugu et al. 1997; Borash et al. 1998) and ammonia (Borash et al. 1998). The evolution of these traits was accompanied, as in the case of the *K*-populations, by a reduced food to biomass conversion efficiency (Joshi and Mueller 1996). CU preadult development time was similar to the UU populations when assayed at low larval density, but lower than the UU populations when assayed at high larval density (Borash and Ho 2001). Urea and ammonia tolerance of the *r*-and *K*-populations were not assayed, while competitive ability of the CU and UU populations was not assayed. The results from these two sets of studies led to what might be termed the canonical view of adaptation to larval crowding in *Drosophila*: the evolution of greater competitive ability was mediated by the evolution of higher feeding rate and greater tolerance to nitrogenous waste, at the cost of efficiency of food utilization (Mueller 1997; Joshi et al. 2001; Prasad and Joshi 2003; Mueller 2009; Mueller and Cabral 2012).

A major element of this canonical view that developed was that increased larval feeding rate is a strong correlate of pre-adult competitive ability in *Drosophila*. Populations subjected to direct selection for increased larval feeding rate also evolved to be better competitors (Burnet et al. 1977), whereas populations selected for rapid pre-adult development evolved both reduced larval feeding rate and reduced larval competitive ability (Prasad et al. 2001; Shakarad et al. 2005; Rajamani et al. 2006). Moreover, populations selected for increased resistance to hymenopteran parasitoids also evolved both reduced larval feeding rate and reduced larval competitive ability (Fellowes et al. 1998, 1999). The suggestion that increased larval feeding rate has a fitness cost unless larval densities are high (Joshi and Mueller 1996) was supported by the observation that larval feeding rates rapidly reverted to control levels in CU populations that were maintained at moderate densities (Joshi et al. 2003). In addition, there also appears to be a trade-off trade-off between urea/ammonia tolerance and larval feeding rate. Populations selected for greater urea and ammonia tolerance, respectively, exhibited the correlated evolution of reduced larval feeding rate (Borash et al. 2000) and larval foraging path length (Mueller et al. 2005), whereas populations selected for greater larval urea tolerance did not exhibit higher pre-adult survivorship than controls at high larval density (Shiotsugu et al. 1997). Overall, results from multiple studies suggest that larval feeding rate and foraging path length are positively correlated (Joshi and Mueller 1988, 1996; Sokolowski et al. 1997; Borash et al. 2000; Prasad et al. 2001; Mueller et al. 2005). Consequently, it appears that the evolution of competitive ability in *Drosophila* depends upon the outcome of a balance between mutually antagonistic traits like increased larval feeding and forgaing behaviour, greater tolerance to nitrogenous wastes, and a reduced efficiency of conversion of food to biomass. Supporting this view, it has been found that the CU populations evolved a development time based polymorphism for two of these traits: compared to controls, offspring of early eclosing flies in a crowded culture showed higher feeding rates, whereas offspring of late eclosing flies showed greater urea/ammonia tolerance (Borash et al. 1998).

Recent studies in our laboratory on species of *Drosophila* other than *D. melanogaster*, however, suggest that there are alternative routes to the evolution of adaptation to larval crowding that do not involve a concomitant increase in larval feeding rate. Populations of *D. ananassae* and *D. nasuta nasuta* subjected to larval crowding under a slightly different maintenance regime than the CU populations evolved greater pre-adult survivorship and larval competitive ability than controls when assayed at high larval density, but without the evolution of increased larval feeding rate, foraging path length and tolerance to nitrogenous wastes, contrary to what was seen earlier in the *K*-and CU populations (Nagarajan et al. 2014). The selected populations of both these species evolved a shorter pre-adult development time, when assayed at both low and high larval densities, and also showed a reduced minimum critical feeding time, the minimum time duration of larval feeding required to sustain successful pupation and eclosion subsequently (Nagarajan et al. 2014). Thus, results from the *D. ananassae* and *D. n. nasuta* crowding adapted populations clearly show that the evolution of greater competitive ability does not necessarily involve the evolution of higher larval feeding rate or foraging path length or tolerance to increasing urea and ammonia in the food medium.

Nagarajan et al. (2014) discussed three possible reasons for the differing phenotypic correlates of increased competitive ability between the studies on *D. ananassae* and *D. n. nasuta* and the earlier studies on *D. melanogaster*. (a) Species-specific differences in the genetic architecture of traits correlated with competitive ability. (b) Differences in genetic architecture between the relatively recently wild-caught populations of *D. ananassae* and *D. n. nasuta* and the long-term laboratory populations of *D. melanogaster* used in the earlier studies, due to very different durations of rearing under domestication. (c) Differences in the details of the laboratory ecology of how larval crowding was implemented between the studies on *D. ananassae* and *D. n. nasuta* and the earlier studies on *D. melanogaster*. The main difference between the maintenance regime used in the *D. melanogaster* studies and that used in the studies on *D. ananassae* and *D. n. nasuta* was in the number of eggs per unit volume of food and the total amount of food used for larval rearing in the crowding-adapted populations (described in detail by Nagarajan et al. 2014). The rearing densities in the *D. ananassae* and *D. n. nasuta* crowded populations were 550-600 eggs in 1.5 mL food, and 350-400 eggs in 2 mL of food, respectively, per vial. In the *K*-and CU populations of *D. melanogaster*, egg densities were not explicitly controlled but were generally higher than those used in the *D. ananassae* and *D. n. nasuta* studies, but with a larger total volume of food, too. It is, therefore, possible that the time course of food depletion and nitrogenous waste build-up in the *D. ananassae* and *D. n. nasuta* crowded cultures is different from that in the *K*-and CU populations. It has been shown theoretically that optimal feeding rates are likely to decline as the concentration of nitrogenous waste in the food increases (Mueller et al. 2005; Mueller and Barter 2015). Thus, at least in principle, it is possible that the optimal feeding rates in the *D. ananassae* and *D. n. nasuta* crowded populations were actually less than they were for the *K*-and CU populations, and that is why increased feeding rates did not evolve in the former (Nagarajan et al. 2014).

In this paper, we describe results from a study of adaptation to larval crowding in long-term laboratory *D. melanogaster* populations that are descendents of the UU control populations used in one of the earlier studies (Mueller et al. 1993; Joshi and Mueller 1996), but are maintained at a density of about 600 eggs in 1.5 mL of food per vial, similar to the maintenance regime of the *D. ananassae* and *D. n. nasuta* crowded populations of Nagarajan et al. (2014). We show that these *D. melanogaster* populations evolve a suite of traits very similar to that seen in the *D. ananassae* and *D. n. nasuta* populations of Nagarajan et al. (2014) and, unlike the earlier studies on *D. melanogaster*, increased competitive ability in these populations is not accompanied by greater larval feeding rate or a reduced efficiency of food utilization. Consequently, these results allow us to rule out genetic architecture differences as a cause for the discrepancies between the studies of Nagarajan et al. (2014) and the earlier *D. melanogaster* studies, especially on the CU populations. It seems clear, therefore, that differences in egg density and total volume of food used in the crowding selection regime can mediate the evolution of increased competitive ability via very different suites of phenotypes.

## Materials and methods

### Study populations

We used eight laboratory populations of *D. melanogaster* for this selection experiment: four populations selected for adaptation to larval crowding (**<u>M</u>**elanogaster **<u>C</u>**rowded as larvae and **<u>U</u>**ncrowded as adults: MCU-1.4), and four ancestral control populations (**<u>M</u>**elanogaster **<u>B</u>**aseline: MB-1.4), whose derivation and ancestral history are summarized in Figure 1. Both the MCU and MB populations were derived from the MGB populations, which were created by mixing the four JB populations (first described by Sheeba et al. 1998), themselves direct descendents of four of the UU populations (Mueller et al. 1993; Joshi and Mueller 1996), and then splitting the four-way hybrid population into five independent replicates, the MGBs (Figure 1).

**Figure 1.**
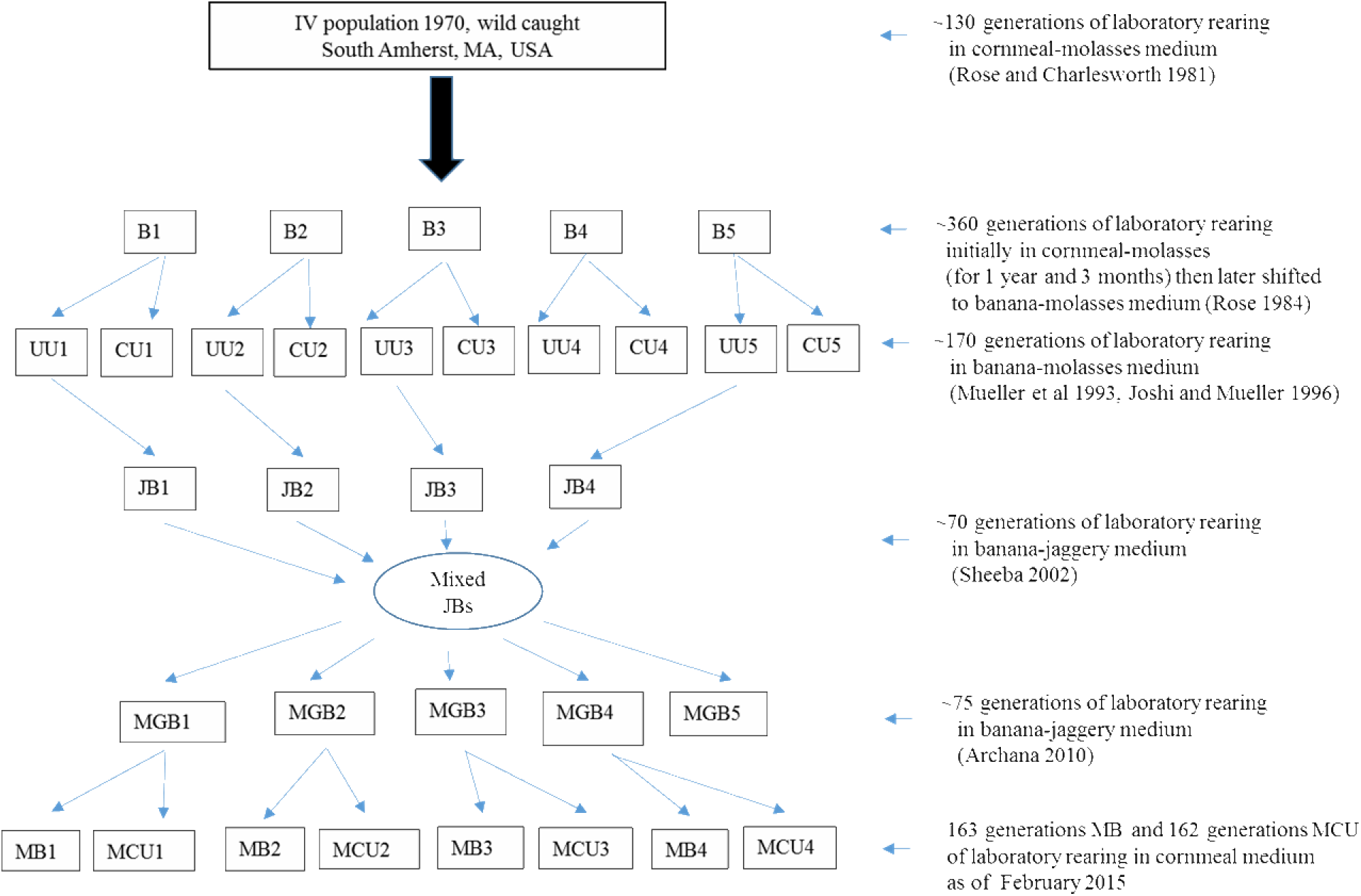
Schematic showing the ancestry of the populations discussed in this study.

The MB populations are maintained at 25 ± 1°C, ~90% relative humidity and constant light on a 21-day discrete generation cycle on cornmeal medium (Table 1). The egg density is kept moderate by collecting 70 ± 10 eggs per vial (9.5 cm height, 2.4 cm inner diameter) in 6 mL of food. Forty such vials are set up per population. On the 11th day after egg collection, the eclosed flies (~75-80% flies eclose by this day) are transferred into Plexiglas cages (25 × 20 × 15 cm^3^). Thus, each population consists of over 2000 breeding adults per generation. In the cage, food plates are changed on the 12^th^, 14^th^ and 17^th^ day after egg collection. On 18th day after egg collection, each cage is provided with a food plate containing excess supplementary live yeast-acetic acid paste. Three days later (ie. on day 21 post egg collection), fresh food plates are provided and the flies are allowed to oviposit for 18 h, and eggs from these plates are transferred into vials to initiate the next generation.

**Table 1.** Composition of cornmeal medium (1 L). For a 1 L cornmeal medium cook, all the ingredients are weighed and added together to 1 L of water. The mixture is then allowed to homogenize on heat, with continuous stirring, after which 120 mL of extra water is added to maintain consistency. The mixture is then pressure cooked for 20 minutes. The prepared medium is then cooled to 60°C, after which preservatives (1g methyl-p-hydroxybenzoate dissolved in 10 ml of ethanol and 1 mL propionic acid) are added.

| Ingredients | Amount |
| --- | --- |
| Cornmeal | 100 g |
| Yeast | 40 g |
| Sugar | 40 g |
| Agar | 12 g |
| Activated charcoal | 0.5 g |
| Water | 1 L |

The MCU populations are maintained in a manner identical to the MB populations except that the rearing density is about 600 eggs in 1.5 mL of food, and that, once eclosions begin, eclosing flies are collected into cages from the rearing vials daily until day 18 post egg collection, a period of roughly 8 days. Initially, the MCU populations were reared at 800 eggs in 1.5 mL of food for a few generations; the density was subsequently reduced to avoid population size crashes. Initially, 24 vials used to be set up per replicate population, but after generation 135 of MCU selection, the number of vials was reduced to 15 per population due to increase in overall egg to adult survivorship over the course of selection. The number of breeding adults is over 2000 per population, as in the MB controls. Each MCU population was derived from one MB population and, hence, MB-*i* and MCU-*i* (*i* = 1.4) are related and were therefore treated as constituting the *i*th block, representing ancestry, in the statistical analyses. The salient features of MB and MCU maintenance, together with that of the CU populations of Joshi and Mueller (1996) are summarized in Table 2 for ease of comparison.

**Table 2.** Similarities and differences in the maintenance regimes of the MB, MCU and CU populations.

| **Population** | **MB** | **MCU** | **CU** |
| --- | --- | --- | --- |
| Larval density | ~70 | ~600 | ~1200-1500 |
| Food volume | 6 mL | 1.5 mL | ~5 mL |
| Type of food | Cornmeal | Cornmeal | Banana-molasses |
| Type of culture vial | 8 dram (2.4 cm diameter × 9.5 cm height) | 8 dram (2.4 cm diameter × 9.5 cm height) | 6 dram (2.2 cm diameter × 8.4 cm height) |
| Fly collection method | Eclosing adults were collected into cages on day 11 from egg collection | Eclosing adults were collected everyday into cages once eclosions began till day 18 post egg collection | Eclosing adults were collected everyday into fresh food vials once eclosions began and kept at a low density of 60-80 adults/vial till day 18 post egg collection |
| Fly collection duration | 1 day | Over 7 days | Over 9 days |
| Yeast supplement duration | 3 days | 3 days | 3 days |
| Generation cycle | 21 day | 21 day | 21 day |

We used two marked mutant populations of *D. melanogaster* as common competitors for the MCU and MB populations in assays of egg to adult survivorship in the presence of a competitor population. One was a white eye population (WE), obtained from spontaneous mutations in one of our wild type JB populations. At the time of assay, the WE population had been maintained in the laboratory for about 90 generations on a 21-day discrete generation cycle on banana-jaggery medium, with all other rearing conditions identical to the MB populations. The second population was an orange eye (OE) mutant population obtained from spontaneous mutations which occurred in the WE population. At the time of assay, the OE population had been maintained for over 48 generations under conditions identical to the MB populations, on cornmeal medium.

### Standardization of populations

Prior to any assay, both control and selected populations were subjected to common (control) rearing conditions for one full generation to eliminate any non-genetic parental effects. About 70 ± 10 eggs were collected per vial in 6 mL of food and 40 such vials were set up per population. On the 11th day after egg collection, the eclosing flies (henceforth referred to as standardized flies) were transferred to cages. Eggs for assays were obtained from these standardized flies over a 14 h period after three days of being provided with food along with a supplement of live yeast-acetic acid paste. All assays were conducted at 25 ± 1°C, under constant light and ~90% relative humidity.

### Egg to adult survivorship at low and high density

Egg to adult survivorship of the MB and MCU populations was assayed at low and high larval density, in the absence (monotypic culture) and presence (bitypic culture) of a marked competitor strain, respectively, at different time points in the course of MCU selection. Egg to adult survivorship at high larval density is the primary trait expected to be under direct selection in the MCU populations exposed to high larval crowding every generation.

#### Generation 30

After 30 generations of MCU selection, eggs laid by standardized flies were placed into vials containing 1.5 mL of food at a density of either 70 or 800 per vial. Eight such vials were set up for each replicate MB and MCU population at each density in monotypic cultures. Eight such vials for each MB and MCU population at each density were also set up in bitypic cultures, with the WE population as a common competitor. For the bitypic cultures, eggs laid by standardized MCU, MB or WE flies were collected and placed into vials containing 1.5 mL of food at a density of either 70 (35 MCU or MB eggs and 35 WE eggs) or 800 (400 MCU or MB eggs and 400 WE eggs) per vial. The number of wild type and mutant (for bitypic cultures) flies eclosing in each vial was recorded and used to calculate egg to adult survivorship in monotypic and bitypic cultures. Vials were checked for eclosion twice a day once eclosions began, till there were no eclosions for four consecutive days.

#### Generation 82

After 82 generations of MCU selection, eggs laid by standardized flies were placed into vials containing 1.5 mL of food at a density of either 60 (ten vials per population) or 600 (six vials per population) per vial. The number of flies eclosing in each vial was recorded and used to calculate egg to adult survivorship in monotypic cultures. Vials were checked for eclosion twice a day once eclosions began, till there were no eclosions for four consecutive days.

#### Generation 112

After 112 generations of MCU selection, eggs laid by standardized flies were placed into vials containing 1.5 mL of food at a density of either 70 (six vials per population) or 600 (five vials per population) per vial. Each vial had half the eggs from an MB or MCU populations and half the eggs from the mutant OE population. The number of wild type and mutant flies eclosing in each vial was recorded and used to calculate egg to adult survivorship in bitypic cultures. Vials were checked for eclosion twice a day once eclosions began, till there were no eclosions for four consecutive days.

### Egg to adult development time at low and high density

After 82 generations of MCU selection, pre-adult development time was assayed at two densities: 60 or 600 eggs per vial with 1.5 mL of food. Eggs from standardized MB and MCU flies were dispensed into each vial using a moistened paintbrush. Eight such vials were set up per replicate population. After the pupae darkened, vials were checked every six hours and the number of eclosing flies recorded. These checks continued till there were no eclosions for four consecutive days.

### Minimum critical feeding time and dry weight after minimum feeding

In this study, conducted at generation 106 of MCU selection, the minimum duration of feeding on yeast required for larvae to successfully complete development was assayed. Eggs for the experiment were collected from the standardized flies which were provided with yeast paste for three days prior to egg collection. Eggs were collected in a narrow time window to synchronize the age of eggs. Eggs from each population were separated into 20 batches of 100-110 eggs each. Each batch was spread on a non-nutritive agar Petri dish (9.5 cm diameter) for hatching. Freshly hatched first instar larvae were subsequently transferred to Petri dishes containing a thin layer of non-nutritive agar overlaid with 37.5% suspension of yeast. Twenty such plates were set up per population, each containing approximately 80 larvae. Larvae were removed from the yeast plates at 3 h intervals between 62 to 80 h from egg collection. At each time point, 90 larvae were removed randomly from these plates, washed, dried on a soft tissue, and then moved to non-nutritive agar vials and their survivorship till eclosion recorded. At each time point, two batches of 15 larvae each per population were also collected, dried in a hot air oven at 70°C for 36 h, and then weighed using a Sartorius (CP 225D) fine balance.

### Dry weight of freshly eclosed adults

After 106 generations of MCU selection, eggs of approximately identical age were harvested over a one hour period from the standardized MB and MCU flies and dispensed onto Petri dishes containing non-nutritive agar at a density of 110 eggs per Petri dish. Eighteen such Petri dishes were set up per population. Twenty two hours later, the freshly hatched first instar larvae from each Petri dish were transferred, using a moistened paintbrush, to a fresh Petri dish containing agar with a thin layer of 37.5% yeast suspension overlaid. Ten such plates with ~100 first instar larvae were set up per population and left undisturbed to allow pupation to occur. After 196 h had elapsed since egg laying, the pupae were transferred to vials with non-nutritive agar, and eventually eclosing adults were harvested from these vials, frozen and subsequently dried in a hot air oven at 64°C for 36 h and weighed in ten batches each of either five males or five females per population using a Sartorius (CP 225D) fine balance.

### Larval feeding rate

After 37 generations of MCU selection, the feeding rates of MB and MCU third instar larvae were measured at physiologically equalized ages, based on the difference in MB and MCU development time. This was done by collecting eggs from the standardized MCU flies 5 h later than the MB flies. Thus, at the time of assay, MCU larvae were ~68 h from egg lay, whereas MB larvae were ~73 h from egg lay. Following Joshi and Mueller (1996), about a hundred eggs laid over a four hour period were collected from standardized flies and placed into two Petri dishes with non-nutritive agar each for the MB and MCU populations. Twenty-four hours later, twenty-five newly hatched larvae were transferred to a Petri dish containing a thin layer of non-nutritive agar overlaid with 1.5 mL of 37.5% yeast suspension. Four such Petri dishes were set up per population. The larvae were then allowed to feed till they were in the early third instar. At this point, 20 larvae from each population were assayed for feeding rate by placing them individually in a Petri dish containing a thin layer of agar overlaid with a thin layer of 10% yeast suspension. After allowing for a 15 sec acclimation period, feeding rate was measured under a stereozoom microscope as the number of cephalopharyngeal sclerite retractions in a 1 min period. Selected and control populations, matched by the subscripted indices, were assayed together, with one larva from the selected population and one from the control population being assayed alternately.

### Larval foraging path length

After 37 generations of MCU selection, the foraging path length of early third instar larvae was measured. The harvesting of larvae was done as described for the larval feeding rate assay. Due to the development time difference between MB and MCU populations, at the time of assay, MCU larvae were ~68 h from egg lay, whereas MB larvae were ~73 h from egg lay. For the assay, 20 larvae from each population were placed individually in a Petri dish containing a thin layer of agar overlaid with a thin layer of 10 % yeast suspension. After allowing for a 15 s acclimation period, the larvae were allowed to move on the Petri plate surface for one minute. The paths that the larvae traversed in a one minute interval were then traced onto a transparency sheet and later measured to the nearest mm with thread and ruler. Selected and control populations, matched by subscripted indices, were assayed together.

### Pupation height

Pupation height was measured after 34 generations of MCU selection for blocks 1 and 2, and after 37 generations of MCU selection for blocks 3 and 4. Eggs from standardized MCU and MB flies were collected off Petri dishes and exactly 30 eggs were placed in a vial with 6 mL of cornmeal medium. Five such vials were set up per population. Once all the individuals had pupated, the pupation heights were measured following Mueller and Sweet (1986) as the distance from the surface of the medium to the point between the anterior spiracles of the pupae. Any pupae on or touching the surface of the food were given a pupation height of zero.

### Larval nitrogenous waste tolerance

After 30 generations of MCU selection, larval tolerance to urea and ammonia in the MB and MCU populations was assayed. Three different concentrations of urea-0, 14, 18 g/L, and three different concentrations of ammonia-0, 15, 25 g/L were used in this assay. Eggs were collected from standardized flies and exactly 30 eggs per 6 mL of food were placed in each vial with different concentrations of urea or ammonia. Ten such vials were set up for each of these three concentrations per population. The number of flies eclosing in each vial was recorded. Nitrogenous waste tolerance was reflected in the mean egg to adult survivorship at different concentrations of urea or ammonia.

### Statistical analyses

Mixed model analyses of variance (ANOVA) were carried out for all the traits. MCU and MB populations with a common numerical subscript were treated as random blocks, which were crossed with the fixed factor selection regime. Depending upon the assay, other fixed factors were larval density, type of culture (monotypic or biypic) for egg to adult survivorship, pre-adult life-stage for instar and pupal duration, urea or ammonia concentration in the food for urea and ammonia tolerance, and larval feeding duration or survivorship for the assays on minimum critical feeding time and dry weight at eclosion after feeding for different time durations. The egg to adult survivorship data were arcsine-square root transformed before ANOVA. Population means were used for testing significance of fixed factors and interactions in all analyses, which were implemented using Statistica for Windows rel. 5.0 B, (Stat Soft 1995). Multiple comparisons were done using Tukey’s HSD test.

## Results

### Egg to adult survivorship at low and high density

In the set of assays done on monotypic and bitypic cultures at low and high density at different times during ongoing MCU selection, the overall trend was for egg to adult survivorship to be higher at low compared to high density, and for MCU populations to have higher egg to adult survivorship than MB controls, with the differences being more pronounced in bitypic cultures with a marked mutant competitor strain (Figure 2, Table 3). At generation 30 of MCU selection, egg to adult survivorship of the MCU populations was significantly higher than MB populations only in bitypic cultures, whereas the effect of density was significant in both monotypic and bitypic cultures (Table 3). Later on in the course of selection, egg to adult survivorship was significantly higher than controls in MCU populations, even in monotypic cultures at generation 82 (Table 3). At generation 112, in bitypic cultures, there was clear evidence for greater competitive ability of the MCU populations, as there was a significant selection regime × density interaction, in addition to significant main effects of selection regime and density (Table 3). At high density, the superiority of MCU populations over MB controls was far greater in the bitypic cultures, in competition with the mutant OE strain than in the montypic cultures (Figure 2b). A similar trend could also be seen in the generation 30 high density survivorship data (Figure 2a), but the interaction between selection regime and density was not significant (Table 3). There was no suggestion of an *r*-*K* trade-off manifesting in reduced survivorship of MCU populations at low density, compared to the MB controls.

**Table 3.**
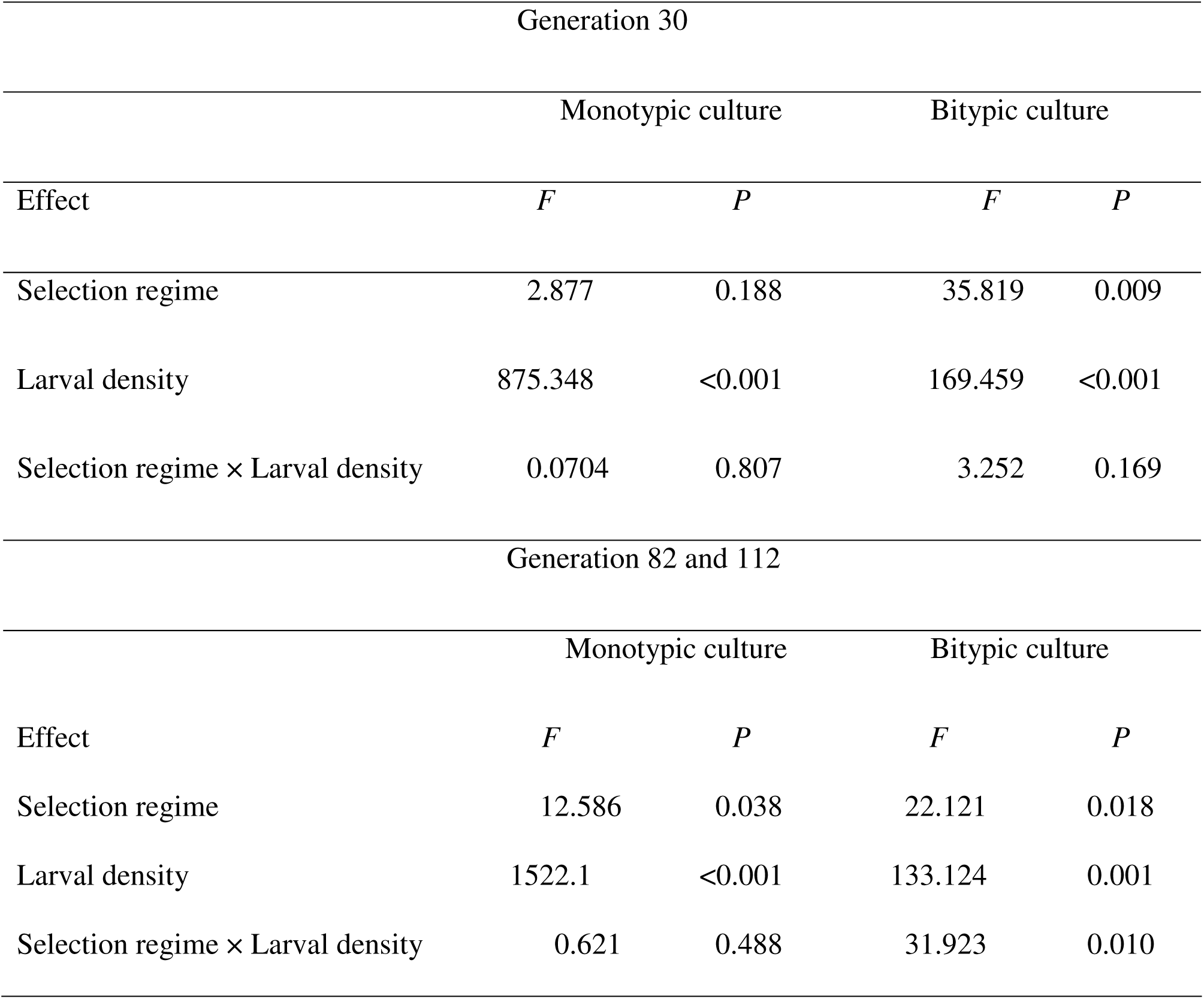
Results of ANOVA on arcsin-squareroot transformed mean egg-to-adult survivorship of MB and MCU populations when competed in monotypic culture and when competed against a common competitor (bitypic culture) at generations 30, 82 and 112 of selection. In this design, the random factor (block) plus any random interactions are not tested for significance and are therefore omitted from the table.

| Generation 30 |  |  |  |  |
| --- | --- | --- | --- | --- |
|  | Monotypic culture |  | Bitypic culture |  |
| Effect | *F* | *P* | *F* | *P* |
| Selection regime | 2.877 | 0.188 | 35.819 | 0.009 |
| Larval density | 875.348 | <0.001 | 169.459 | <0.001 |
| Selection regime × Larval density | 0.0704 | 0.807 | 3.252 | 0.169 |
| Generation 82 and 112 |  |  |  |  |
|  | Monotypic culture |  | Bitypic culture |  |
| Effect | *F* | *P* | *F* | *P* |
| Selection regime | 12.586 | 0.038 | 22.121 | 0.018 |
| Larval density | 1522.1 | <0.001 | 133.124 | 0.001 |
| Selection regime × Larval density | 0.621 | 0.488 | 31.923 | 0.010 |

**Figure 2.**
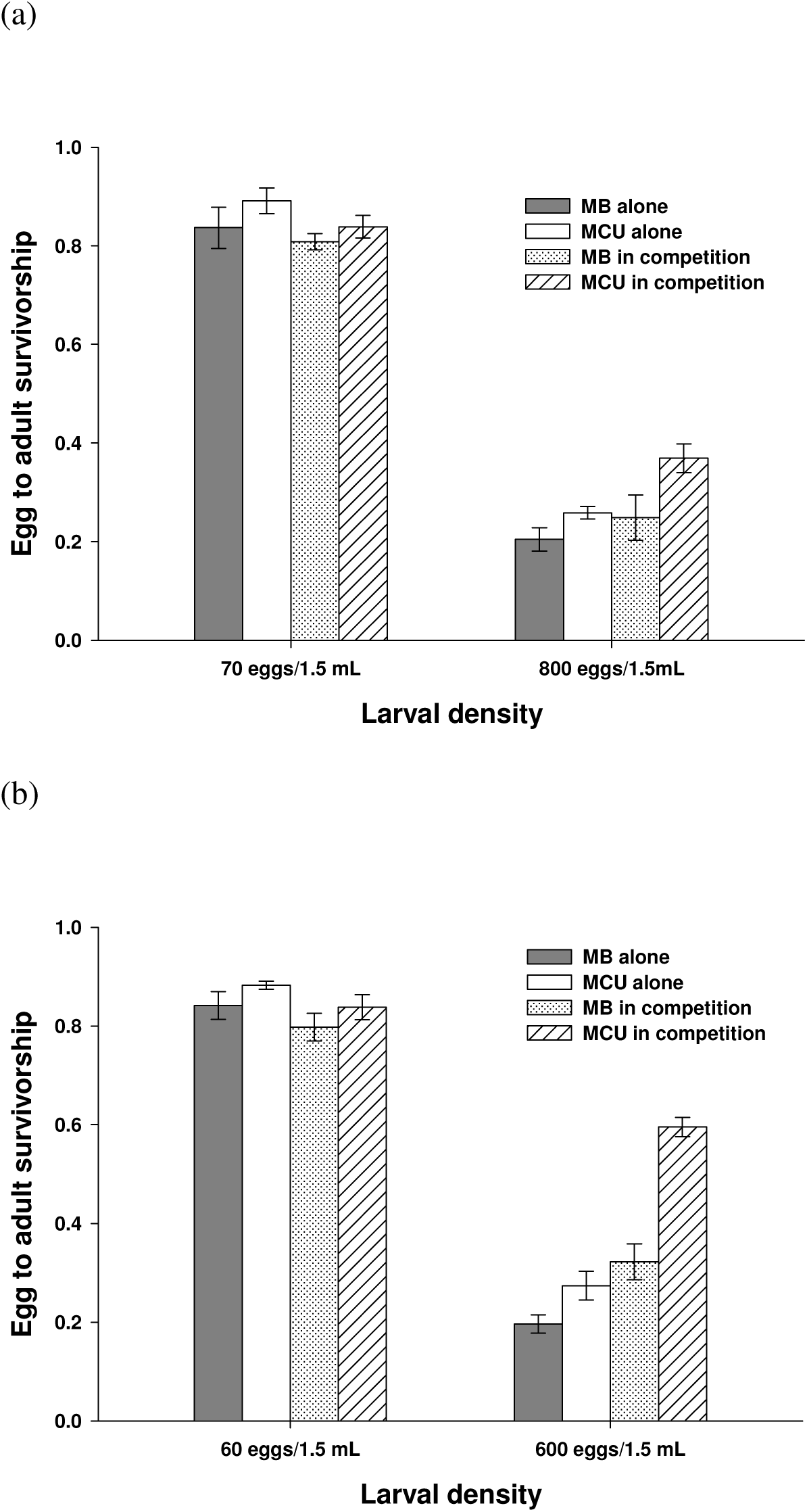
Larval competitive ability after generation (a) 30 of selection (b) 82 and 112 of selection. The graphs represent mean egg-to-adult survivorship of MB and MCU populations in the competition assay (either in a monotypic culture or in bitypic culture with a common competitor) at moderate and high larval density. Error bars are the standard errors around the means of the four replicate populations.

### Egg to adult development time at low and high density

The ANOVA revealed a significant main effect of selection regime (Table 4), with the MCU populations developing significantly faster than the MB controls at both low and high density (Figure 3). The magnitude of the development time difference was about 10h in males and 15h in females at both low and high densities (Figure 3), driving a significant selection regime × sex interaction (Table 4). The other significant ANOVA effects were those of density (slower development at high density) and the sex × density interaction (Table 4). Overall, females developed significantly faster than males at low density whereas the male-female difference was not significant at high density (Tukey’s HSD test at *P* = 0.05 level of significance). In the MCU populations, females developed faster than males at both densities, whereas in the MB populations, females had similar development time as males at low density, but took about 3.5 h longer than males to complete development at high density (Figure 3). None of these differences, however, were significant by Tukey's HSD test.

**Figure 3.**
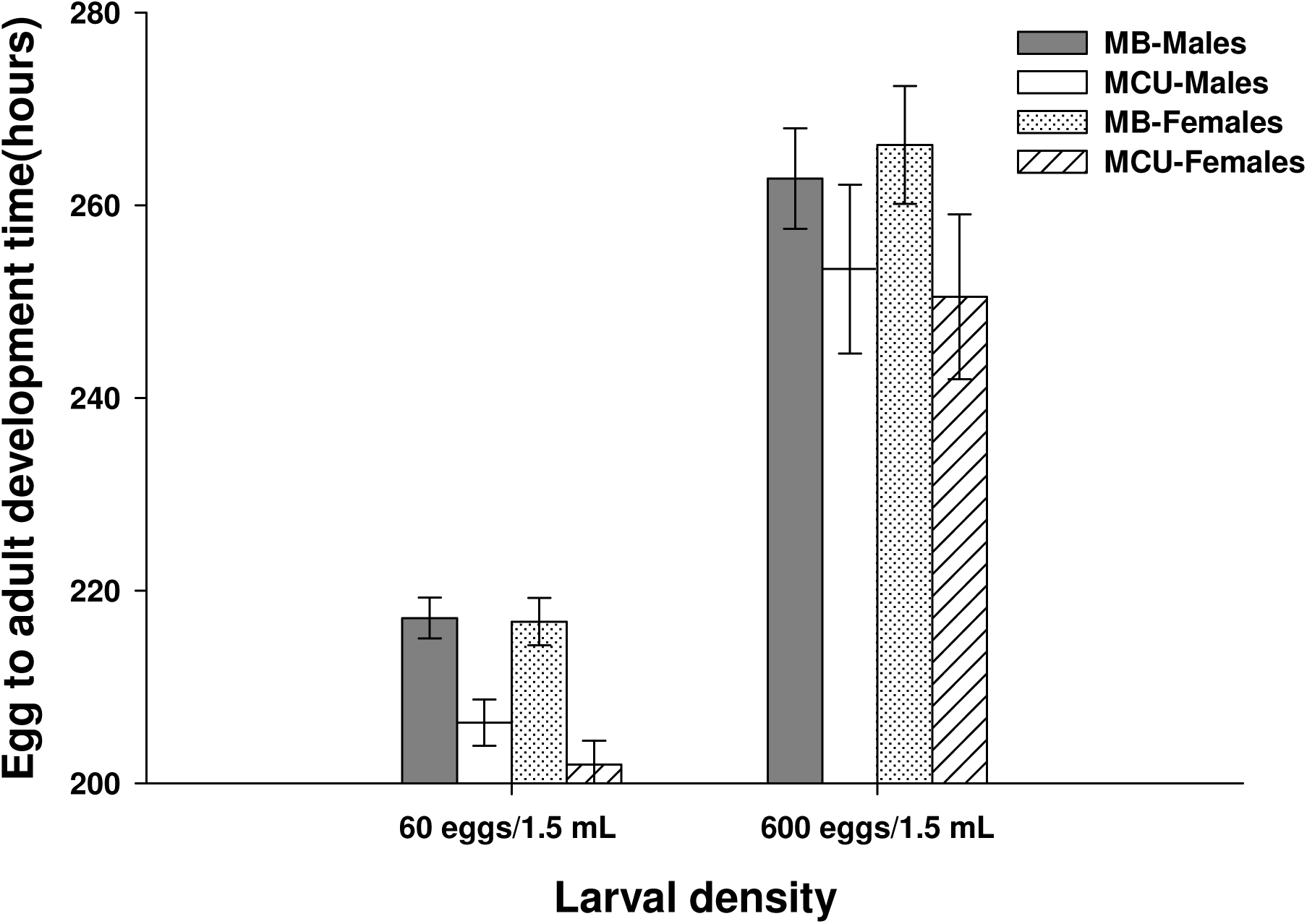
Egg to adult development time of MB and MCU populations after 82 generations of selection at moderate and high larval density. Error bars are the standard errors around the means of the four replicate populations.

**Table 4.**
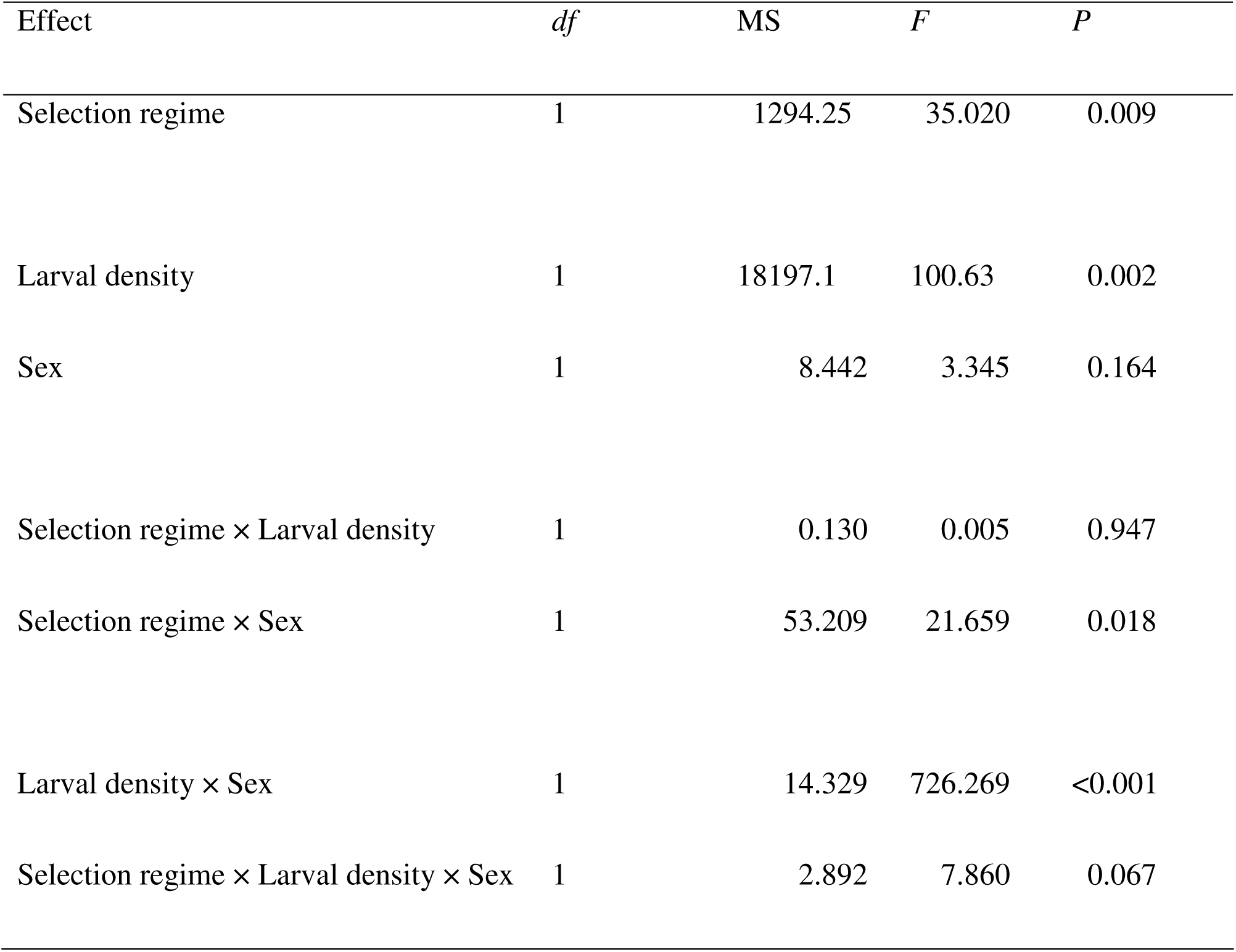
Results of ANOVA on mean egg to adult development time assayed at low and high larval density after 82 generations of MCU selection, with selection regime, larval density and sex as fixed factors, and block as a random factor. In this design, the random factor (block) plus any random interactions are not tested for significance and are therefore omitted from the table.

| Effect | *df* | MS | *F* | *P* |
| --- | --- | --- | --- | --- |
| Selection regime | 1 | 1294.25 | 35.020 | 0.009 |
| Larval density | 1 | 18197.1 | 100.63 | 0.002 |
| Sex | 1 | 8.442 | 3.345 | 0.164 |
| Selection regime × Larval density | 1 | 0.130 | 0.005 | 0.947 |
| Selection regime × Sex | 1 | 53.209 | 21.659 | 0.018 |
| Larval density × Sex | 1 | 14.329 | 726.269 | <0.001 |
| Selection regime × Larval density × Sex | 1 | 2.892 | 7.860 | 0.067 |

### Minimum critical feeding time and dry weight after minimum feeding

The survivorship to adulthood of larvae allowed to feed for different durations of time after egg collection (62, 65, 68, 71, 74, 77 or 80 h) tended to increase with feeding duration for both MB and MCU populations (Figure 4a, Table 5: significant main effect of feeding duration). At each time point after 65 h from egg collection, however, MCU larvae showed significantly greater survivorship to adulthood than the MB controls (Figure 4a, Table 5: significant main effect of selection regime). There was no significant selection regime × feeding duration interaction (Table 5).

**Table 5.** Results of ANOVA on mean survivorship to eclosion of larvae from the MB and MCU populations on non-nutritive agar, after having been allowed to feed for different durations of time. In this design, the random factor (block) plus any random interactions are not tested for significance and are therefore omitted from the table.

| Effect | *df* | MS | *F* | *P* |
| --- | --- | --- | --- | --- |
| Selection regime | 1 | 0.315 | 297.848 | < 0.001 |
| Feeding duration | 6 | 0.538 | 29.609 | < 0.001 |
| Selection regime × Feeding duration | 6 | 0.024 | 2.365 | 0.073 |

**Figure 4.**
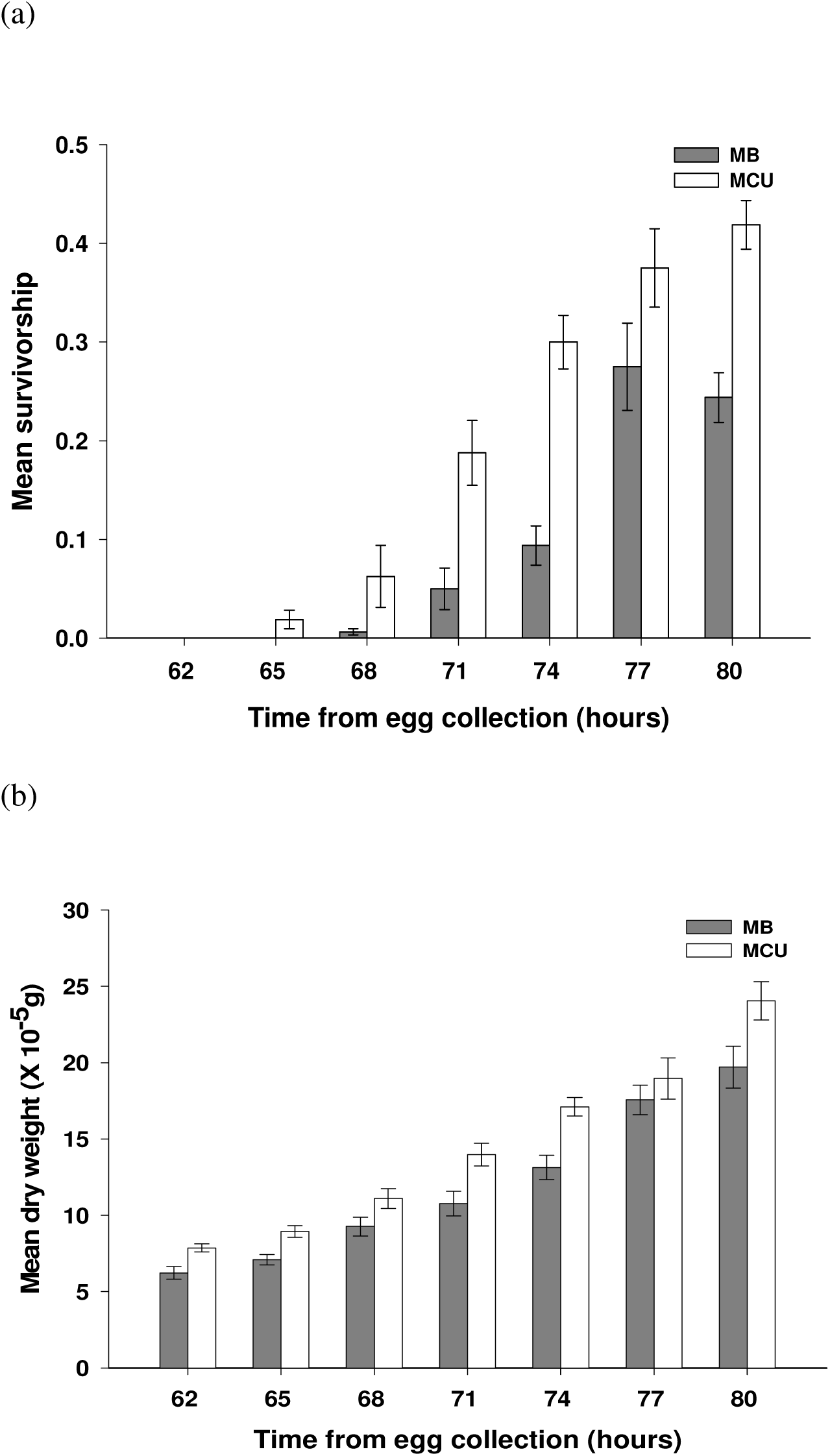
(a) Mean survivorship to eclosion of larvae when removed from food at different time points from egg lay and transferred to non-nutritive agar plates. (b) Mean dry weight of individual larvae when measured at different time points from egg collection. Error bars are the standard errors around the means of the four replicate populations.

The dry weight of larvae allowed to feed for different durations of time after egg collection (62, 65, 68, 71, 74, 77 or 80 h) also tended to increase with feeding duration for both MB and MCU populations (Figure 4b, Table 6: significant main effect of feeding duration). At each time point, the MCU larvae were significantly heavier (except at 77 h) than the MB controls (Figure 4b, Table 6: significant main effect of selection regime). There was no significant selection regime × feeding duration interaction (Table 6). The time points covered (62-80 h from egg collection) correspond roughly to mid-late second instar through approximately middle third instar.

**Table 6.** Results of ANOVA on mean dry weight of larvae at different stages of larval development, in a span of 62 to 80 h from egg collection in MB and MCU populations, with selection and time as fixed factors and block as a random factor. In this design, the random factor (block) plus any random interactions are not tested for significance and are therefore omitted from the table.

| Effect | df | MS | *F* | *P* |
| --- | --- | --- | --- | --- |
| Selection regime | 1 | 95.298 | 8832.264 | < 0.001 |
| Feeding duration | 6 | 238.881 | 58.033 | < 0.001 |
| Selection regime × Feeding duration | 6 | 2.944 | 58.033 | 0.430 |

### Dry weight of freshly eclosed adults

The ANOVA on dry weight of freshly eclosed adults revealed significant main effects of selection regime and sex (Table 7). Females were significantly heavier than males, and both MCU males (mean ± s.e. = 30.39 × 10^−5^ ± 0.40 × 10^-5^ g) and females (mean ± s.e. = 40.82 × 10^−5^ ± 0.40 × 10^−5^ g) were significantly lighter than MB males (mean ± s.e. = 30.96 × 10^−5^ ± 0.42 × 10^−5^ g) and females (mean ± s.e. = 42.07 × 10^−5^ ± 0.95 × 10^−5^ g), respectively.

**Table 7.**
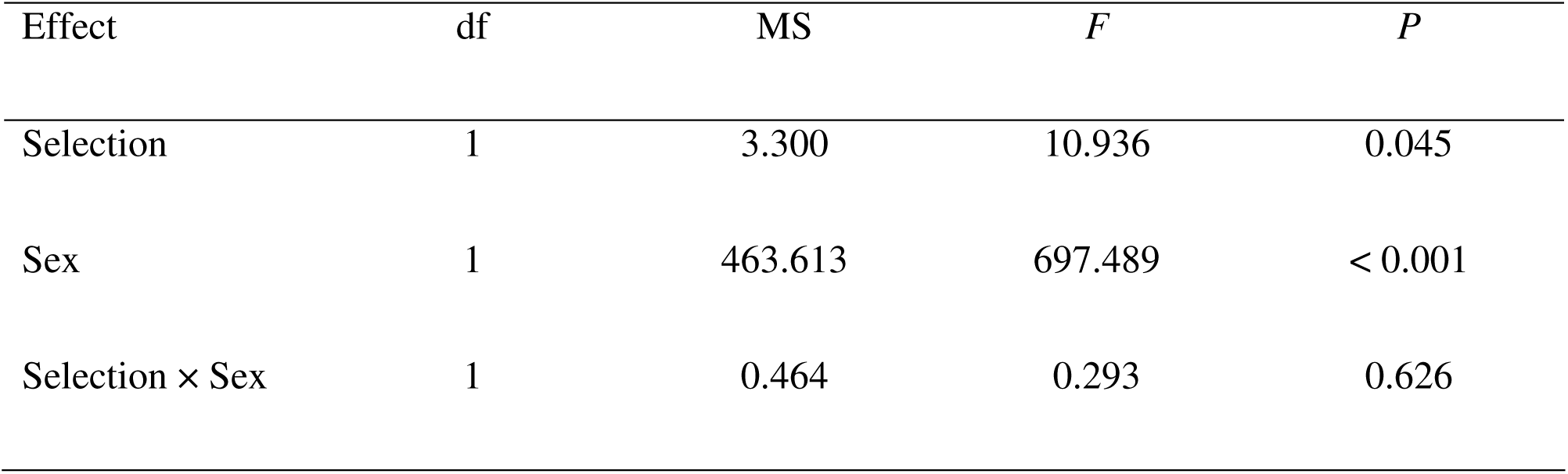
Results of ANOVA on mean dry weight at eclosion of MB and MCU populations, with selection and sex as fixed factors and block as a random factor for both analyses. In this design, the random factor (block) plus any random interactions are not tested for significance and are therefore omitted from the table.

### Larval behaviours

The MB and MCU populations did not differ in any of the three larval behaviours assayed. Larval feeding rates in the MCU (mean ± s.e. = 108.8 ± 1.23 bites per min) and MB (mean ± s.e. = 108.2 ± 1.36 bites per min) populations did not differ significantly (*F*_1,3_ = 0.51; *P* = 0.53). Larval foraging path lengths also did not differ significantly (*F*_1,3_ = 1.93; *P* = 0.26) between the MCU (mean ± s.e. = 3.65 ± 0.036 cm) and MB (mean ± s.e. = 3.83 ± 0.033 cm) populations. Pupation heights, too, did not differ significantly (*F*_1,3_ = 1.44; *P* = 0.32) between the MCU (mean ± s.e. = 1.18 ± 0.11 cm) and MB (mean ± s.e. = 1.36 ± 0.088 cm) populations.

### Larval nitrogenous waste tolerance

There was no evidence for any difference between the MCU and MB populations in tolerance to either ammonia (Figure 5a) or urea (Figure 5b). The only significant ANOVA effect on survivorship on both ammonia and urea was that of waste concentration (Table 8), and in the case of both ammonia and urea, survivorship of both MCU and MB populations declined with increasing concentration (Fig. 5).

**Table 8.** Results of three-way ANOVA on egg to adult survivorship of MB and MCU populations in food containing different concentrations of ammonia and urea after 30 generations of selection. Selection and concentrations were fixed factors and block was a random factor.

| Effect | *df* | MS | *F* | *P* |
| --- | --- | --- | --- | --- |
| Ammonia |  |  |  |  |
| Selection | 1 | 0.003 | 3.82 | 0.145 |
| Concentration | 2 | 1.17 | 627.73 | <0.001 |
| Selection × Concentration | 2 | 0.0013 | 1.64 | 0.27 |
| Urea |  |  |  |  |
| Selection | 1 | 0.012 | 4.83 | 0.115 |
| Concentration | 2 | 1.99 | 89.45 | <0.001 |
| Selection × Concentration | 2 | 0.007 | 2.69 | 0.146 |

**Figure 5.**
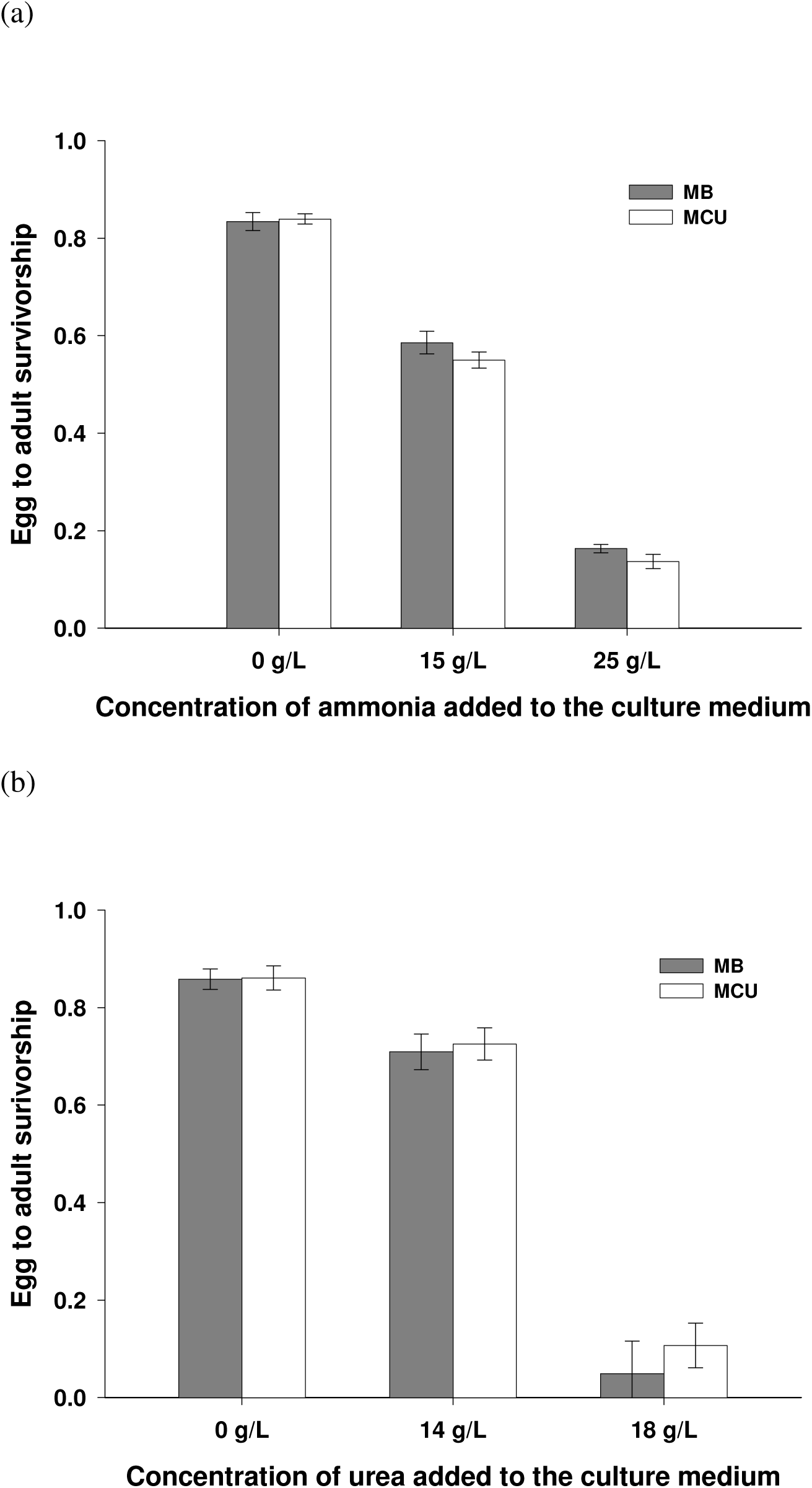
Mean egg-to-adult survivorship of MB and MCU populations in three different concentrations of (a) ammonia, and (b) urea, after 30 generations of selection. Error bars represent the standard errors around the means of the four replicate populations.

## Discussion

The present study was motivated by the earlier observation that crowding adapted populations of *D. ananassae* and *D. n. nasuta* evolved greater competitive ability and pre-adult survivorship at high density primarily through a combination of reduced duration of the larval stage, faster attainment of minimum critical size for pupation and greater efficiency of food conversion to biomass (Nagarajan et al. 2014). This was in contrast to the generally accepted view that the evolution of greater competitive ability in fruitflies was primarily mediated by the evolution of increased larval feeding rate, foraging path length and tolerance to nitrogenous wastes, at the cost of reduced efficiency of food conversion to biomass, based on earlier studies with *D. melanogaster* (Mueller 1997; Prasad and Joshi 2003; Mueller 2009). We decided to ascertain whether the differences in the phenotypic correlates of greater competitive ability between these two sets of studies were due to the different species used, or differences in the manner in which larval crowding was imposed, by examining the direct and correlated responses to selection for adaptation to larval crowding in populations of *D. melanogaster* related to those used in the earlier studies (UU populations, described by Joshi and Mueller 1996), but maintained under a rearing protocol similar to that used in the studies on *D. ananassae* and *D. n. nasuta* by Nagarajan et al. (2014).

Our results clearly show that the MCU populations of *D. melanogaster* have adapted to crowded larval conditions by evolving traits very similar to those seen in the *D. ananassae* and *D. n. nasuta* crowding adapted populations by Nagarajan et al. (2014), and very different from those seen in the CU populations of *D. melanogaster* (Mueller 1997; Prasad and Joshi 2003; Mueller 2009). Specifically, like the crowding adapted *D. ananassae* and *D. n. nasuta* populations (Nagarajan et al. 2014), the MCU populations appear to have evolved greater competitive ability (Figure 2b) largely through reduced development time (Figure 3), and a reduction in the minimum larval feeding duration needed to be able to successfully complete pre-adult development (Figure 4a), perhaps accompanied by a reduction in the minimal critical size for pupation. Moreover, unlike the CU populations of *D. melanogaster* studied earlier (Mueller 1997; Prasad and Joshi 2003; Mueller 2009), the MCU populations do not differ from MB controls in larval feeding rate, foraging path length, pupation height or nitrogenous waste tolerance (Figure 5).

Despite the broad similarities in correlated responses to selection for adapting to high larval density, however, there are some differences between the results reported here and the results on similar selected populations of *D. ananassae* and *D. n. nasuta* reported by Nagarajan et al. (2014). Our MCU populations did not show any evidence of an *r*-*K* trade-off (MCU survivorship was higher than controls at both low and high larval density: Figure 2), whereas the crowding adapted *D. ananassae* and *D. n. nasuta* populations showed a trend of lower survivorship than controls when assayed at low larval densities (Nagarajan et al. 2014). Moreover, a population dynamics study on the *D. ananassae* populations suggested that *r* was actually lower in the crowding adapted populations, as compared to their controls (Dey et al. 2012). In the MCU populations, the development time reduction, relative to controls, was not magnified at high larval density (Figure 3), whereas the development time difference between the crowding adapted and control *D. ananassae* and *D. n. nasuta* populations almost doubled when assayed at high larval density (Nagarajan et al. 2014). The crowding adapted *D. ananassae* and *D. n. nasuta* populations evolved greater pupation height than their respective controls (Nagarajan et al. 2014), but the MCU pupation height did not differ from controls. At this point, we do not have clear explanations for these differences in some of the correlated responses to selection across the three sets of populations.

In the crowding adapted *D. ananassae* populations, there was evidence for the evolution of reduced minimum critical time for feeding without a reduction in the minimum critical size for successfully completing subsequent development to adulthood (Archana 2010; Nagarajan et al. 2014). Although survival in the *D. ananassae* selected populations was greater than controls after different durations of feeding, the dry weights of eclosing flies that had fed as larvae for different durations of time did not differ between selected and control populations. In contrast, larvae from the MCU populations exhibited a time advantage (head-start) over their controls of about 6 h in terms of survivorship after feeding for different durations (Figure 4a), and a similar advantage of about 3 h for larval dry weight after feeding for different durations (Figure 4b). This suggests a reduction not only in minimum critical feeding time, but also in minimum critical size, in the MCU populations, relative to controls. Interestingly, till about mid third instar, MCU larvae are heavier than their control counterparts (Figure 4b) but they subsequently become lighter than controls during the wandering larva stage (Sarangi 2013) and eventually eclose as lighter adults than their control counterparts (Table 7). This pattern suggests that MCU populations have evolved a greater larval growth rate during second and early to mid third instar, in addition to reduced minimum critical size, and then a further reduction in the late third instar duration, leading them to stop feeding correspondingly earlier than controls, and life-stage-specific development time data support this view (Sarangi 2013).

Despite the few differences between correlated responses to selection shown by the MCU populations on the one hand, and the *D. ananassae* and *D. n. nasuta* crowding adapted populations on the other, as discussed above, the overall similarity in the traits through which all three sets of crowding adapted populations in our laboratory evolved greater larval competitive ability clearly rules out two of the three hypotheses put forward by Nagarajan et al. (2014). The observation that the MCU populations, which share common ancestry with the CU populations of Joshi and Mueller (1996), nevertheless evolve a suite of traits similar to that seen in the crowding adapted *D. ananassae* and *D. n. nasuta* populations rules out differences between species, or between relatively recently wild-caught versus long-term laboratory adapted populations, in genetic architecture of traits relevant to fitness under larval crowding as being the cause for the different responses to selection seen in the earlier studies with *D. melanogaster* (Joshi and Mueller 1996) and the studies of Nagarajan et al. (2014). The remaining possibility, thus, becomes the most likely explanation: that differences in the details of the laboratory ecology of how larval crowding was implemented between the studies on the MCU populations and the crowding adapted populations of *D. ananassae* and *D. n. nasuta* on the one hand, and the earlier studies on *D. melanogaster* on the other, led to the evolution of very different suites of traits conferring increased competitive ability in the two sets of studies.

As noted earlier, the main difference between the maintenance regime used in the earlier *D. melanogaster* studies and that used in this study and the studies on *D. ananassae* and *D. n. nasuta* was in the number of eggs per unit volume of food and the total amount of food used for larval rearing in the crowding-adapted populations (described in detail by Nagarajan et al. 2014). Compared to the three sets of crowding adapted populations used in our laboratory, the CU cultures had both a larger number of eggs and a greater total volume of food. It is, therefore, likely that the time course of food depletion and nitrogenous waste build-up in the MCU, *D. ananassae* and *D. n. nasuta* crowded cultures is different from that in the earlier studied *D. melanogaster* populations. Specifically, in vials with high larval density but only 1.5 ml of food (as in the MCU, *D. ananassae* and *D. n. nasuta* crowded cultures) the food gets used up very fast and probably also accumulates toxic levels of nitrogenous waste quite quickly as the total amount of food available for waste to diffuse into is small even early on. *Drosophila* larvae typically feed within a 1 cm depth from the surface of the food, and in vials with 1.5 ml of food, the height of the food column is less than 1 cm. If food levels are closer to 5-6 ml (as in the CU populations), there is greater amount of food for waste to diffuse into and thus the effective waste concentration experienced by feeding larvae in the 1 cm depth feeding band is likely to be lower than that experienced by larvae in vials with only 1.5 ml of food, especially in the early stages of the crowded culture. At the same time, since feeding is restricted to a 1 cm deep band, effective competition for access to food, especially in early stages of a crowded culture, will be higher in vials with a greater absolute number of larvae (as in the CU populations), because the volume of food in the 1 cm deep feeding band is constant, but the number of larvae feeding in that zone is higher in the CU cultures compared to the MCU, *D. ananassae* and *D. n. nasuta* crowded cultures. As it has been shown theoretically that optimal feeding rates are likely to decline as the concentration of nitrogenous waste in the food increases (Mueller et al. 2005; Mueller and Barter 2015), it is likely that the optimal feeding rates in the 1.5 mL crowded cultures are lower than if there is 5-6 mL food per vial. Additionally, since food runs out very fast over time in crowded vials with only 1.5 mL of food, it might not permit better survival of individuals with a high waste tolerance, unlike the case in crowded vials with 5-6 mL of food.

If the speculation above is correct, then the nature of selection acting on traits affecting competitive ability in *Drosophila* cultures is likely to depend critically on both egg density and total amount of food, specifically the height of the food column. This also suggests that although measures like egg density are often of heuristic value when thinking about adaptation to crowding, ultimately the ecological details of different types of crowded cultures can affect the form of the fitness functions acting on different traits related to competitive ability, thereby mediating the evolution of greater competitive ability via different sets of trait changes. In principle, it should be possible to directly test these predictions both by doing selection experiments at different combinations of egg density and total food volume, as well as by directly examining the effects of different combinations of egg density and total food volume on the phenotypic distributions of traits relevant to fitness under larval crowding in fruitflies.

## Acknowledgments

We thank Larry Mueller for much helpful discussion, D. Ravi Teja, Avani Mital, N. Rajanna and M. Manjesh for help in the laboratory, and N. G. Prasad for very helpful and Industrial Research, Government of India, for financial assistance in the form of Junior and Senior Research Fellowships. S. Dey and M. Sarangi were supported by doctoral fellowships from the Jawaharlal Nehru Centre for Advanced Scientific Research. This work was supported by funds from the Department of Science and Technology, Government of India, to A. Joshi. The preparation of the manuscript was supported in part by a J. C. Bose National Fellowship to A. Joshi. The authors declare no competing interests.

## References

Anderson W. W. and Arnold J. 1983 Density-regulated selection with genotypic interactions. Am. Nat. 121, 649–655.

Archana N. 2010 The genetic architecture of fitness-related traits in populations of three species of Drosophila subjected to selection for adaptation to larval crowding. Ph.D. thesis, Jawaharlal Nehru Centre for Advanced Scientific Research, Bengaluru, India.

Asmussen M. A. 1983 Density-dependent selection incorporating intraspecific competition. II. A diploid model. Genetics 103, 335–350.

Borash D. J., Gibbs A. G., Joshi A. and Mueller L. D. 1998 A genetic polymorphism maintained by natural selection in a temporally varying environment. Am. Nat. 151, 148–156.

Borash D. J., Teótonio H., Rose M. R. and Mueller L. D. 2000 Density-dependent natural selection in *Drosophila*: correlations between feeding rate, development time and viability. J. Evol. Biol. 13, 181–187.

Borash D. J. and Ho G. T. 2001 Patterns of selection: stress resistance and energy storage in density-dependent populations of *Drosophila melanogaster*. J. Insect Physiol. 47, 1349–1356.

Burnet B., Sewell D. and Bos M. 1977 Genetic analysis of larval feeding behaviour in *Drosophila melanogaster*. II. Growth relations and competition between selected lines. Genet. Res. 30, 149–161.

Clarke B. 1972 Density-dependent selection. Am. Nat. 106, 1–13.

Dey S., Bose J. and Joshi A. 2012 Adaptation to larval crowding in *Drosophila ananassae* leads to the evolution of population stability. Ecol. Evol. 2, 941–951.

Fellowes M. D. E., Kraaijeveld A. R. and Godfray H. C. J. 1998 Trade-off associated with selection for increased ability to resist parasitoid attack in *Drosophila melanogaster*. Proc. R. Soc. Lond. B 265, 1553–1558.

Fellowes M. D. E., Kraaijeveld A. R. and Godfray H. C. J. 1999 Association between feeding rate and parasitoid resistance in *Drosophila melanogaster*. Evolution 53, 1302–1305.

Gadgil M. and Bossert, W. H. 1970 Life historical consequences of natural selection. Am. Nat. 104, 1–24.

Joshi A. and Mueller L. D. 1988 Evolution of higher feeding rate in *Drosophila* due to density-dependent natural selection. Evolution 42, 1090–1092.

Joshi A. and Mueller L. D. 1993 Directional and stabilizing density-dependent natural selection for pupation height in *Drosophila melanogaster*. Evolution 47, 176–184.

Joshi A. and Mueller L. D. 1996 Density-dependent natural selection in *Drosophila*: trade-offs between resource acquisition and utilization. Evol. Ecol. 10, 463–474.

Joshi A., Prasad N. G. and Shakarad M. 2001 *K*-selection, *α*-selection, effectiveness and tolerance in competition: density-dependent selection revisited. J. Genet. 80, 63–75.

Joshi A., Castillo R. B. and Mueller L. D. 2003 The contribution of ancestry, chance and past and ongoing selection to adaptive evolution. J. Genet. 82, 147–162.

MacArthur R. H. 1962 Some generalized theorems of natural selection. Proc. Natl. Acad. Sci. USA 48, 1893–1897.

MacArthur R. H. and Wilson E. O. 1967 The theory of island biogeography. Princeton University Press, Princeton, NJ, USA.

Mueller L. D. and Ayala A. J. 1981 Trade-off between *r*-selection and *K*-selection in *Drosophila* populations. Proc. Natl. Acad. Sci. USA 78, 1303–1305.

Mueller L. D. and Sweet V. F. 1986 Density-dependent natural selection in *Drosophila*: evolution of pupation height. Evolution 40, 1354–1356.

Mueller L. D. 1988 Evolution of competitive ability in *Drosophila* by density-dependent natural selection. Proc. Natl. Acad. Sci. USA 85, 4383–4386.

Mueller L. D. 1990 Density-dependent selection does not increase efficiency. Evol. Ecol. 4, 290–297.

Mueller L. D., Graves J. L. and Rose M. R. 1993 Interactions between density-dependent and age-specific selection in *Drosophila melanogaster*. Func. Ecol. 7, 469–479.

Mueller L. D. 1997 Theoretical and empirical examination of density-dependent selection. Annu. Rev. Ecol. Syst. 28, 269–288.

Mueller L. D., Folk D. G., Nguyen N., Nguyen P., Lam P., Rose M. R. and Bradley T. J. 2005 Evolution of larval foraging behavior in *Drosophila* and its effect on growth and metabolic rates. Physiol. Entomol. 30, 262–269.

Mueller L. D. 2009 Fitness, demography, and population dynamics in laboratory experiments. In Experimental evolution: concepts, methods and applications of selection experiments (Garland T. Jr., Rose M. R., eds.), pp. 197–216. University of California Press, Berkeley, CA, USA.

Mueller L. D. and Cabral L. G. 2012 Does phenotypic plasticity for adult size versus food level in *Drosophila melanogaster* evolve in response to adaptation to different rearing densities? Evolution 66, 263–271.

Mueller L. D. and Barter T. T. 2015 A model of the evolution of larval feeding rate in *Drosophila* driven by conflicting energy demands. Genetica (doi: 10.1007/s10709-015-9818-5).

Nagarajan A., Natarajan S. B., Jayaram M., Thammanna A., Chari S., Bose J., Jois S. V. and Joshi A. 2014 Adaptation to larval crowding in *Drosophila ananassae* and *Drosophila nasuta nasuta*: increased larval competitive ability without increased larval feeding rate. BioArxiv (doi: http://dx.doi.org/10.1101/011684).

Pianka E. R. 1970 On *r*-and *K-*selection. Am. Nat. 104, 952–956.

Prasad N. G., Shakarad M., Anitha D., Rajamani M. and Joshi A. 2001 Correlated responses to selection for faster development and early reproduction in *Drosophila*: the evolution of larval traits. Evolution 55, 1363–1372.

Prasad N. G. and Joshi A. 2003 What have two decades of laboratory life-history evolution studies on *Drosophila melanogaster* taught us? J. Genet. 82, 45–76.

Rajamani M., Raghavendra N., Prasad N. G., Archana N., Joshi A. and Shakarad M. 2006 Reduced larval feeding rate is a strong evolutionary correlate of rapid development in *Drosophila melanogaster*. J. Genet. 85, 209–212.

Rose M. R. and Charlesworth B. 1981 Genetics of life-history in *Drosophila melanogaster*. I. Sib analysis of adult females. Genetics 97, 173–186.

Rose M. R. 1984 laboratory evolution of postponed senescence in *Drosophila melanogaster*. Evolution 38, 1004–1010.

Roughgarden J. 1971 Density-dependent natural selection. Ecology 52, 453–468.

Sarangi M. 2013 Preliminary investigations into the causes for alternative routes to the evolution of competitive ability in populations of Drosophila selected for adaptation to larval crowding. M.S. thesis, Jawaharlal Nehru Centre for Advanced Scientific Research, Bengaluru, India.

Shakarad M., Prasad N. G., Gokhale K., Gadagkar V., Rajamani M. and Joshi A. 2005 Faster development does not lead to correlated evolution of greater competitive ability in *Drosophila melanogaster*. Biol. Lett. 1, 91–94.

Sheeba V. 2002 Probing the adaptive significance of circadian rhythms using Drosophila melanogaster. Ph.D. thesis, Jawaharlal Nehru Centre for Advanced Scientific Research, Bengaluru, India.

Sheeba V., Madhyastha N. A. A. and Joshi A. 1998 Oviposition preference for normal versus novel food resources in laboratory populations of *Drosophila melanogaster*. J. Biosci. 23, 93–100.

Shiotsugu J., Leroi A. M., Yashiro H., Rose M. R. and Mueller L. D. 1997 The symmetry of correlated responses in adaptive evolution: an experimental study using *Drosophila*. Evolution 51, 163–172.

Sokolowski M. B., Pereira H. S. and Hughes K. 1997 Evolution of foraging behavior in *Drosophila* by density-dependent selection. Proc. Natl. Acad. Sci. USA 94, 7373–7377.

StatSoft. 1995 Statistica Vol. I: general conventions and statistics 1. StatSoft Inc., Tulsa, OK, USA.

